# Phenotypic scoring of Canola Blackleg severity using machine learning image analysis

**DOI:** 10.1101/2025.07.17.665372

**Authors:** Qiao Hu, Sarah Anderson, Stuart Gardner, Thomas W. Ernst, Chadwick B. Koscielny, Navratan S. Bahia, Charles G. Johnson, Andrew C. Jarvis, Joseph S. Hynek, Nathan Coles, Igor Falak, David R. Charne, Manuel E. Ruidiaz, Julien Linares, Anastasios Mazis, Daniel J. Stanton

**Author notes:** Corresponding author: Daniel J. Stanton, 22220 Highway 16, Ardrossan AB Canada T8E 2L4. Abbreviations: CNN (convolutional neural networks); DL (deep learning); EU (experimental unit); G (genotype); GT (ground truth); BB (bounding box); ML (machine learning); BLUP (best linear unbiased prediction).

## Abstract

Canola blackleg is a fungal disease that causes significant yield loss and plant death of infected canola (*Brassica napus* L., *Brassica rapa L.*, *Brassica juncea L.*) fields worldwide. One of the most effective methods for controlling blackleg is through the cultivation of resistant varieties. Consequently, scoring blackleg disease severity of infected plants is a key metric for identifying and selecting resistant varieties. Traditionally, blackleg severity is scored by expert raters who evaluate disease in stem cross sections using established rating scales and reference images; however, human raters are expensive and inconsistent in their scoring. Here, we introduce a machine learning algorithm based on deep learning models that can score blackleg severity from cross-section images of infected plants. We find that expert ratings are largely inconsistent across raters and across years for the same rater, creating substantial noise in susceptibility ratings. Meanwhile, our trained machine learning model performs more consistently than the median rater while maintaining a similar heritability as expert raters for the blackleg susceptibility trait. This model can be used to standardize blackleg susceptibility scoring across locations and years to improve canola breeding outcomes across affected regions.

**Core Ideas:**

- Canola Blackleg is a fungal disease affecting yield of canola, and accurate scoring of Blackleg severity is important for tracking disease and breeding for resistant varieties.
- The standard practice of utilizing expert raters is expensive, and scores assigned are inconsistent across raters and years.
- Our deep learning model for assigning blackleg severity scores is more accurate than the median expert rater, opening the door for improved breeding of new resistant varieties.

## Introduction

Canola/rapeseed (*Brassica napus* L., *Brassica rapa L.*, *Brassica juncea L.*) is the world’s second-largest oilseed crop (Wanasundara et al., 2017), with Canada contributing around 20% of the world’s production. Due to its high adaptability to diverse climates, particularly mesic temperate regions (Jaime et al., 2018), and its suitability for soil pH values ranging from 5.5-8.3 (Myers, 2018), canola has become an economically important crop for many countries and regions including Canada, Australia, USA, India, China, and the EU. Between 2005 and 2021, canola was the second most lucrative agricultural sector in Canada, contributing $26.7 billion to the country’s GDP (Windfeld & Lhermie, 2022). At the same time, global efforts are underway to increase the farming efficiency of the crop, with the European Union achieving the highest farming efficiency (∼2.76 t/ha) in a 10-year average (2010-2020), demonstrating the importance of the crop as source of oil for biofuel (Zandberg et al., 2022).

The economic value of the species has motivated breeding efforts that focus on developing varieties resilient to climate change and resistant to pathogens, including Canola Blackleg (or Blackleg for short). Blackleg (or phoma stem canker) is an endemic disease of several *Brassica* species grown in North America, Australia, and Europe, and is caused by the fungi *Leptosphaeria maculans* and *Leptosphaeria biglobosa*. Blackleg was first detected in Saskatchewan in 1975 and has since become widespread throughout the province (Gugel & Petrie, 1992). It is a saprophytic disease that persists year to year on infected plant stubble (Naseri, 2006). The airborne spores, called ascospores, infect plants as early as the cotyledon stage, resulting in small lesions on the leaves and sometimes plant death. Plants that are infected later in the season can develop blocked vascular tissue that can girdle the plant, restricting the flows of water and nutrients necessary for complete pod fill. After harvest, infected plant residues remain in the field resulting in conditions for spore dispersion the following season (Zhang XueHua & Fernando, 2018). Efforts to reduce the incidence of the disease include crop rotation, stubble management, fungicides, selection of resistant varieties or a combination of the above (Hwang et al., 2016; West et al., 2001). Varieties with resistance to blackleg, such as Quantum, Q2, Hi-Q and Conquest (Zhang & Fernando, 2017), have been available in Canada since the 1990s (Kutcher et al., 2013); however, blackleg still remains one of the most serious pathogens that threaten canola production, resulting in periodic large-scale economic losses (Bokor et al., 1975; West et al., 2001), reducing seed yield by up to 50% in extreme cases (Gugel & Petrie, 1992).

One of the most effective methods for controlling widespread blackleg outbreaks has been variety selection (Zhang XueHua & Fernando, 2018). Genetic resistance to blackleg is provided in two basic ways: quantitative and qualitative. While quantitative genetic resistance to blackleg provides some protection, major resistance genes such as *RLM3* have been critical to the development of durable blackleg resistant varieties. Evidence suggests blackleg resistance is beginning to breakdown in *RLM3* and other major R-genes (Zhang et al., 2016), meaning breeders will either have fewer tools to develop new resistant varieties or will need to identify and introduce new sources of genetic resistance. A key to identifying new sources of genetic resistance is accurate phenotyping of the disease, in a way that is predictive of potential yield loss.

During the 1980s and 1990s, several scoring methods and scales were proposed as the standard for canola blackleg (Cargeeg & Thurling, 1980; Gugel et al., 1990; Koch et al., 1989; Kutcher, 1990; Newman & Bailey, 1987). Gugel et al. (1990) and Berg et al. (1993) found that blackleg ratings based on internal symptoms like vascular tissue discoloration were more predictive than methods based on external visible phenotypes (Gugel et al., 1990; Van den Berg et al., 1993). Today, blackleg susceptibility is most often quantified by measuring the amount of discoloration from a stem cross-section of an infected plant, with many countries (such as Canada) requiring a blackleg score for all registered canola varieties planted.

The current standard for disease scoring, established by the Canola Council of Canada, uses a 0 – 5 scale, with classes of disease ranging from no visible disease in the cross-section of the stem base (0) to plants without any living tissue or lodged (5). Other scales have been proposed (1 – 6 scale by Aubertot et al., 2004 or empirical 1– 9 scale by breeders), with all of them following the same protocols for visually assessing the severity of the disease before they are converted to the CCC standard scale. However, the phenotype attributes can vary from plant to plant, and it requires highly trained individuals to accurately score the disease. Scorers rely on sketches and pictures of the symptoms as a reference to standardize their disease ratings. In addition, phenotyping stem discoloration is labor intensive, including the need to identify diseased plants, dissect them at the proper spot, and accurately score the diseased area on a very short period during the growing season. To alleviate some of the above issues, a software was developed to help train new scorers to the 1– 6 scoring scale (Aubertot et al., 2005) utilizing image analysis as one way to enhance the accuracy and precision of blackleg scoring, but it was considered too time-consuming based on the technology available at the time (Aubertot et al., 2004).

Today, image analysis is increasingly used to monitor crop growth and development, assess plant architecture and traits, and detect stress in plants. In many crops, image analysis can also be used to score diseases utilizing unmanned aerial vehicles (Kaivosoja et al., 2021; Wu et al., 2019) and proximal phenotyping methods (DeChant et al., 2017; Ngugi et al., 2020) to assess external disease symptoms. Particularly, thanks to the breakthrough of deep learning (DL) in 2010s (Krizhevsky, 2012), advances in machine learning methods for computer vision tasks, like using convolutional neural networks (CNNs) (He, 2016; Ronneberger et al., 2015; Simonyan, 2015; Szegedy, 2015), have resulted in many validated disease scoring techniques that can replace or augment manual scoring in breeding programs (Arsenovic et al., 2019; Lu et al., 2021; Saleem et al., 2020). Disease scoring based on images can greatly improve data collection protocols, increase the scoring accuracy, and democratize scoring so that researchers without specialized skills in pathology can collect disease data (Bousset et al., 2018; Wu et al., 2019).

Herein, we demonstrate the development and assessment of a deep learning algorithm to accurately score canola Blackleg in a time and labor efficient way while standardizing the disease scoring.

## Materials and Methods

### Field Experiment Setup

Canola was planted for phenotyping trials near Corteva research stations throughout the canola growing regions of Canada in 2018 and 2019. Single row plots containing a range of varieties including both blackleg susceptible and resistant genotypes of canola were planted, and four replicate plots were planted for each genotype. Blackleg inoculum of *L. maculans* genotypes AvrLm2-3-4-5-6-7-9-11, AvrLm3-4-5-6-7-9-11, and AvrLm2-3-5-6-9-11 were prepared for the field based on previously published methods (Fraser et al., 2020). Both grain inoculum and pycnidiospore suspensions were applied to the single row plots. Grain inoculum was applied by hand directly post-emergence at a rate of 25g per single row plot, while pycnidiospore suspensions were foliar applied to plants once the first true leaf was visible with approximately 1L of inoculum (1 × 10^6 spores mL−1) sprayed per row at about 30 lbs psi.

Between the stages of seed set and maturity, the optimal time for swathing, plants were harvested for evaluation of Blackleg disease severity. A target of 25-30 plants were pulled from the middle of each plot, avoiding any plants with thin stems (<3mm) or that appeared to be infected by a disease other than blackleg. Plants were gathered into bundles by human operators and were loaded into a stationary imaging machine to collect image samples. Plants were oriented with the root crown interface aligned with the cutting blade of the machine. The conveyor includes an air bladder to ensure alignment of the stem for cutting by the internal blade. The stem and cross section were then imaged in a photometry box and then conveyed further to discard out the end of the platform. Images of the cut stems were sampled inside a darkened box, with two 5MP Basler ace acA2500-20gc cameras collecting images both perpendicular (for a cross-section view) and parallel (for a profile view of the infected area) to the stem. Both views of images are in RGB color space and have the size of 2592 by 2048 pixel. Due to the machine assembly and camera calibration accuracy of each year, minor differences between images of these two years in image profiles such as color balance and general cross-section placements are observed. Examples of images in each year are in Supplemental Figure 1.

### Manual Scoring

To develop and evaluate the deep learning model, the ground truth (GT) of blackleg infestation level on collected samples was generated via human experts viewing the images and assigning a rating. The rating is a 1 –9 scale system where 1 represents the most severe with 100% of internal tissue diseased, 9 represents the least severe disease rating with no lesions, and intermediate scores of 2-8 correspond to >75%, 51-75%, 50%, 25-50%, 25%, <25% and <10% of internal tissue diseased, respectively. In addition, a score of “Unsure” is included for samples in any cases where a rating cannot be generated from the image acquisition through the decision-making processes. Unsure ratings may indicate that the image quality is too low to score, the cross-section is too difficult to read, the cross-section is missing in the image or corrupted, for example, blocked by mud or other structures, or the cut of the cross-section is too low to the root or too high, hence the infected area is not imaged. Examples of 1–9 scale plus Unsure are in Figure 1. When an expert rates samples, both the cross-section and profile images may be present to rate. The cross-section is the primary reference for the rating, but the profile view is supplementary and can help to differentiate between severe-infection cases, particularly where a score of 1 is assigned to plants with a dead cross-section and dead stem while a score of 2 assigned when the stem is still green despite a dead cross-section. For this reason, when a cross-section is corrupted or impaired, a sample is viewed as Unsure; and when only a profile is corrupted or impaired, a sample is still rated with experts’ best effort. The GT rating was generated using the median of all raters’ ratings. For samples where more than half of the experts rate a sample as Unsure, then its GT is Unsure; otherwise, Unsure ratings are disregarded when calculating the GT. If the median is a float due to the presence of Unsure, the average is rounded to the closest integer rating as the GT. For 2019 samples with GT values that are low (<=3) or high (>=8), 4 of 5 expert raters evaluated both the cross-section and profile images again, and the total 9 ratings (5 original and 4 new) were used to determine the GT.

**Figure 1:**
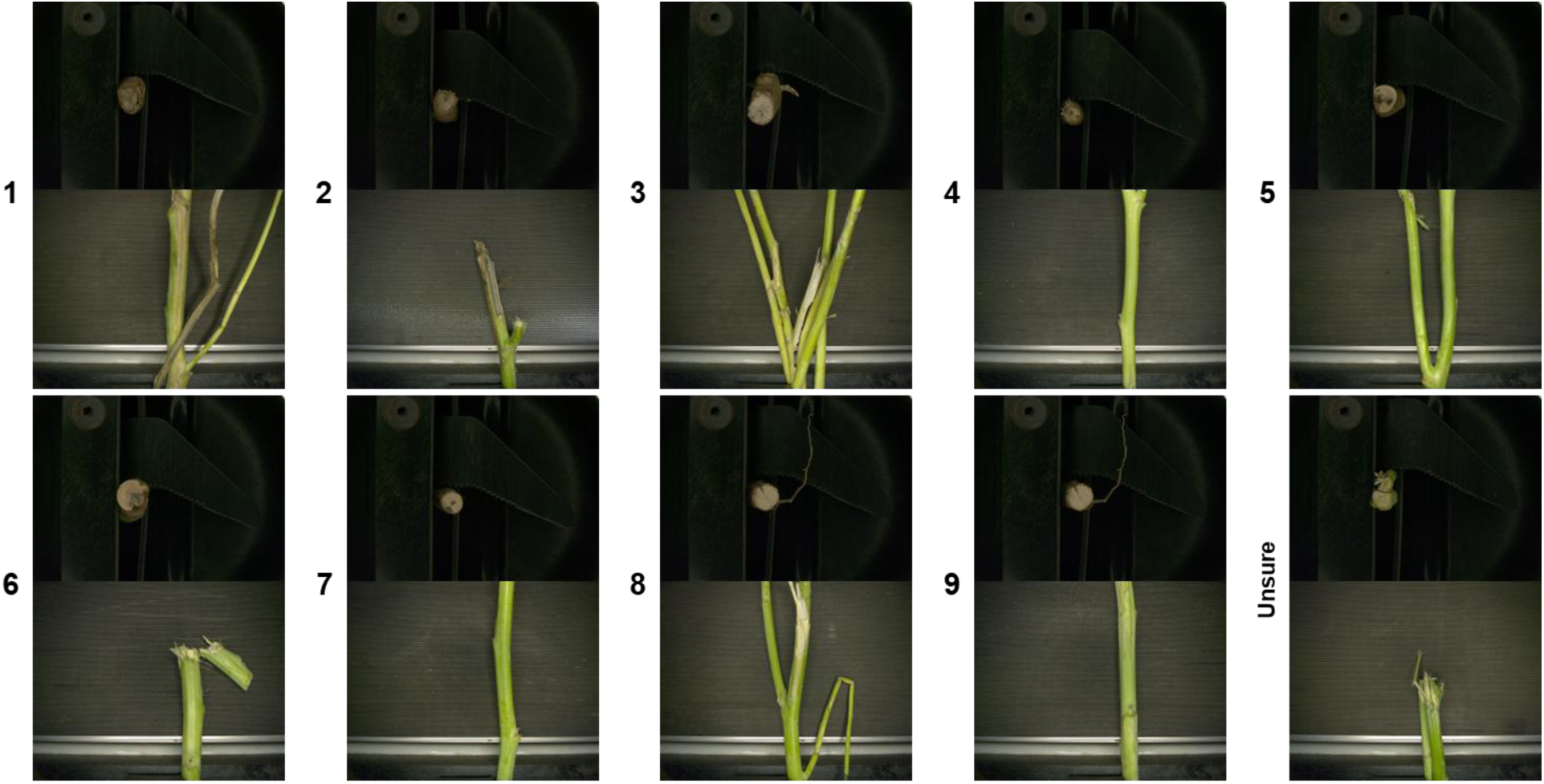
Examples of canola samples with blackleg scores from 1-9 plus Unsure. The Unsure example shows the case where two cross-sections exist, indicating the cut is too high.

### Image Processing Architecture

The demonstrated image analysis method is a two-stage method. A diagram of the method is in Figure 2.

**Figure 2:**
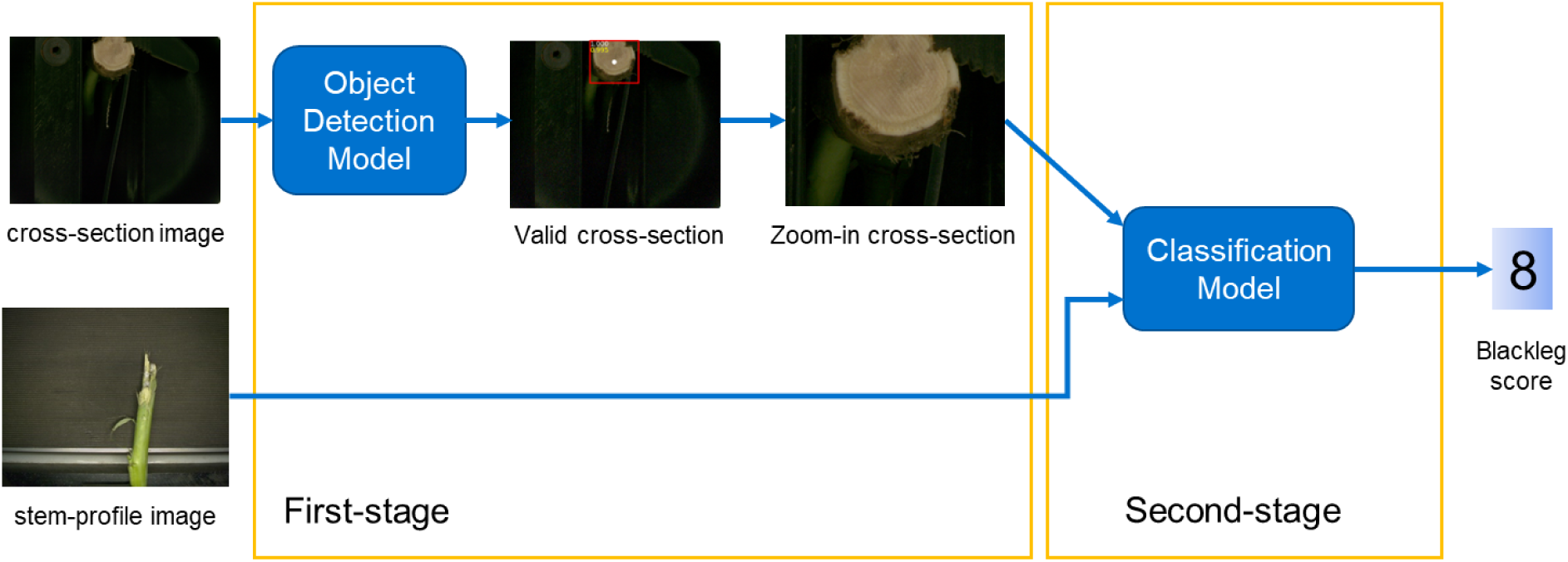
Diagram of the two-stage DL-based system. The first-stage object detection model accepts as input a cross-section image and outputs a zoomed-in cross-section image. The second-stage classification model accepts as input a pair of images, and the output is the blackleg severity score.

The first-stage model is an object detection (Girshick, 2015; Girshick, 2014; Liu, 2016) deep learning (DL) model which serves multiple purposes. The first objective is to detect if there is a valid cross-section that is feasible to rate. The second is to crop the image against the cross-section so the image is zoomed-in, standardized in dimension, and centered when feeding the image to the second-stage model, facilitating the model’s differentiation on the infected area. For these purposes and for computation efficiency, a lightweight YOLO V3 model (Redmon, 2018) is used here. When training a YOLO model, training data of cross-section images and the rectangular bounding box (BB) coordinate of the cross-section as the GT are fed to the YOLO model. The model yields BB coordinates as its output, and the difference between the model’s yield and the GT coordinate is minimized as the objective of the training process. Once the training is complete, with input of an image, the model outputs one or multiple BBs with their coordinates and each of the BB includes a cross-section. In addition, the model generates for each BB a confidence score of a 0 to 1, indicating whether it contains a valid cross-section or not. The confidence score is used to determine one BB by selecting the highest score when there are multiple BBs, and also as an indicator if there is a valid cross-section or not. This is because one image should have either one valid canola sample, or no valid sample (Unsure) because of any errors mentioned earlier. Hence, a threshold for the confidence is used as the model’s parameter so that when the confidence is below the threshold, the cross-section is invalid. In this case, the second stage is skipped, and the sample is determined as Unsure. Otherwise, the image is cropped using the center of the BB with a fixed size and sent to the second stage for further rating.

For the second stage, a customized CNN model based on the ResNet architecture (He, 2016) was developed to produce the severity rating. The ResNet is a supervised image classification model that, after training, yields a class for a given image. The classes are the severity ratings from 1 to 9. In order to utilize both the cross-section and the profile image of the stem, a customization is made to the model that both images are stacked together as the input of the model, and the model’s first layer is extended from 3 channels (for originally a single RGB image) to 6 channels (for the stacked two RGB images). Once the model is trained, it accepts as input the cropped cross-section generated by the first-stage model and the profile image, and it generates a score between 1 –9 along with a confidence associated with the score. A threshold is also used here to assign a low confidence score as Unsure. Combining the two models, the system can generate a score including Unsure for any given pairs of images acquired via our imaging system. This two-stage architecture is designed because of the field-of-view concern for the cross-section and the need to use both view images, over a single object detection DL model with a modification to handle the two-view input. The field-of-view concern is that the cross-section image needs to have a large field-of-view because the placement of the sample is not precisely controlled. This causes an issue that in the image, the cross-section could be small and at a relatively random location (see the various cross-section examples in Figures 3-4). Therefore, both stages are required for confident scoring of blackleg severity. Model Training

**Figure 3:**
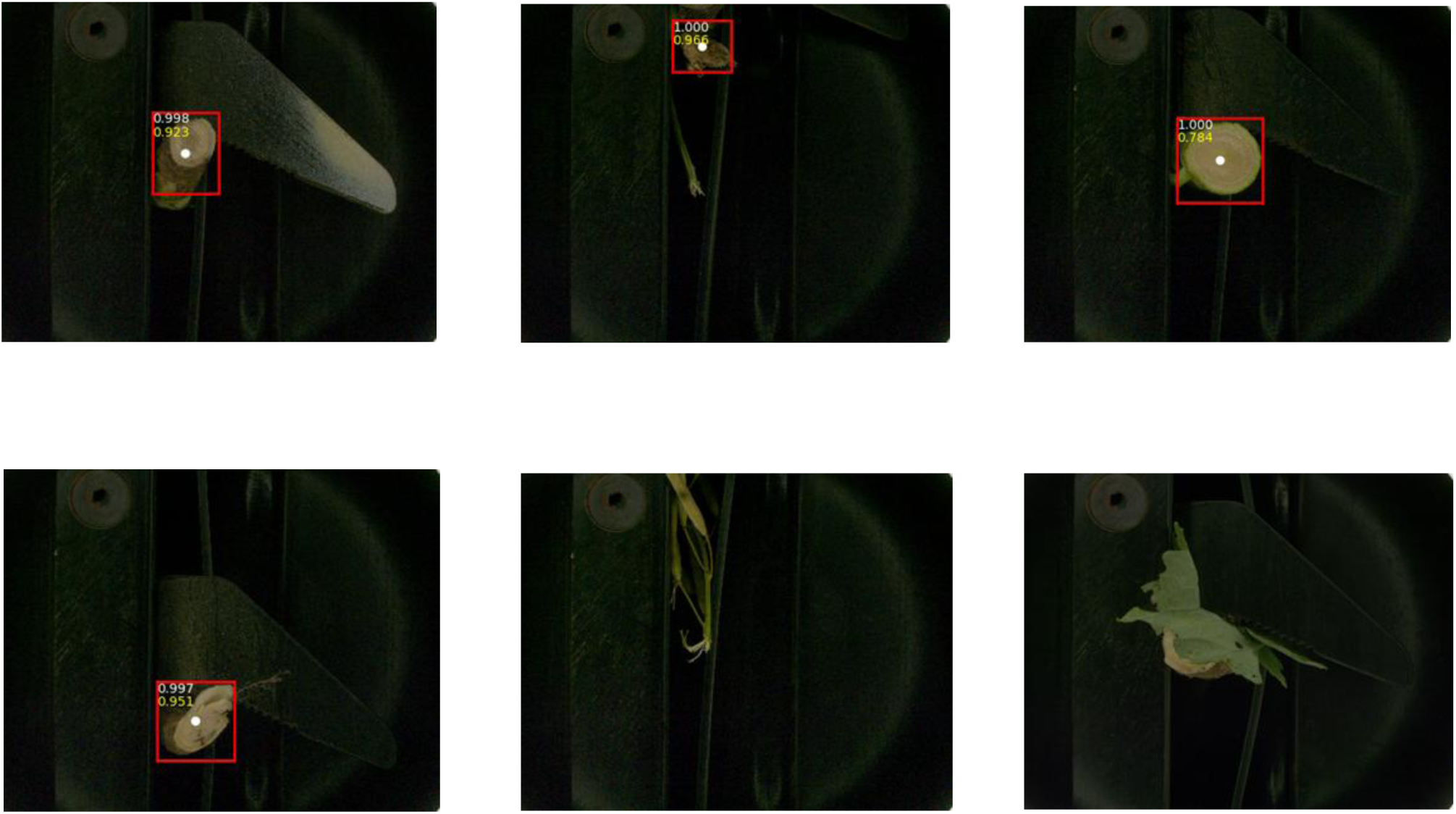
examples of correct decisions by the first-stage model. The red rectangle indicates a bounding box and the below number in yellow indicates the confidence score. The last two examples are Unsure cases and the model outputs no bounding box and returns Unsure label as the result.

**Figure 4:**
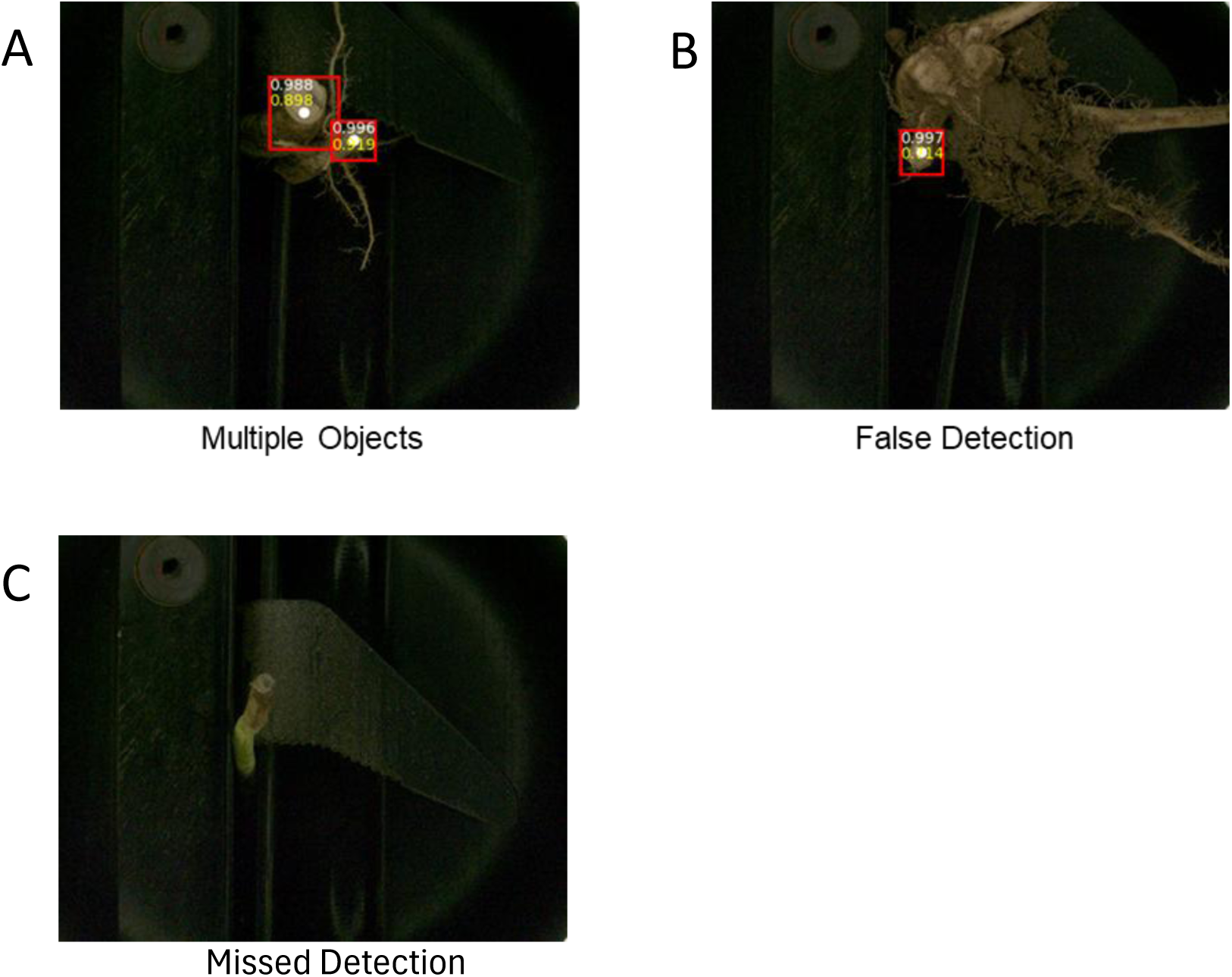
Examples of errors of various types resulting in an Unsure rating (A-B) or missed detection (C). (A) Example of sample resulting from multiple invalid cross-sections presents due to high-cut, where the model detects both. (B) Example of sample due to low-cut and with mud blockage, where the model detects one cross-section falsely. (C) Example of sample showing a possible cross-section but the model fails to detect it.

Since both models are supervised DL models, images collected for 2018 and 2019 are split into training and testing sets to train and evaluate the models individually. For both models, each pair of the cross-section image and the profile image is considered the independent sample.

For the first-stage model, samples from 2018 and only the cross-section images are used to develop the model and split randomly into a training and testing set with a 36-64% ratio, respectively. All images were manually annotated by humans with a rectangle BB to tightly include the cross-section of the stem when it is visible. Images of Unsure are not annotated with a BB but are still included in the training or testing set as negative samples. For the model, the official Pytorch implementation (Redmon, 2018) of the YOLO V3 is used, and a transfer learning method is used by fine-tuning (Donahue, 2014; Oquab, 2014; Razavian, 2014) on a pre-trained published YOLO model provided along the implementation. In this case, the YOLOv3-416 is used so that the input size of the image is 416 by 416, and hence all images are rescaled to this size when fed to the model. Data augmentation is used during the training where images are randomly flipped vertically and/or horizontally, and they are randomly cropped so that the center of the image is a random shift from the center of the BB. As training parameters, the standard optimizer Adam (Kingma, 2017) is used, the batch size is 64, and the epoch is 100. Once the model is trained, the confidence threshold is empirically set by obtaining the best results on the training set so that every image can be correctly called for whether there is a ratable cross-section or not. In this case and also for simplicity, the threshold is set as 0.5, although thresholds of 0.2, 0.25, and 0.3 were also tested before determining the optimal value. Prior to inputting into the second-stage model, all images are cropped at the center of the detected BB with a fixed 1298x1298 size as the input of the second stage.

For the second stage model, similar to the first-stage model, fine-tuning was based on a pre-trained published model implemented and developed in Torchvision (Ansel et al., 2024; Contributors, 2017; He, 2015). For training and testing this model, samples from both 2018 and 2019 are used. They are randomly split with the stratification to the score distribution. Samples of 2018 are split with an 85%-15% ratio for the training and testing set, and samples of 2019 are split with a 1%-99% ratio. Then the training set is the combination of 2018 training set and 2019 training set. For samples where no BB is produced by the first-stage model, these samples are skipped in the training. However, for any sample where the profile image is missing or of low-quality, it is still included in the training or testing set as long as a BB is detected. In a particular case where a profile image is missing, a fixed profile image without a stem profile is used as a substitute. Because the model is modified so that the first kernel’s channel is extended from 3 to 6 for training on the stacked cross-section and profile images, the weight of the first layer of the pretrained model is duplicated to match the expansion of the 3 to 6 channels. As training parameters, Adam optimizer is also used, the batch size is 64, and the epoch is 100. The best model is selected with the lowest loss within the 100 epoch.

### Evaluation of Heritability and BLUPs

Statistical analysis of the phenotypic data was conducted on the subset of testing data from 2019 from one location (Ardrossan). Analysis was performed using ASReml software (Butler, 2023) on the experimental plot unit (EU) level. This one location was chosen since it is the only 2019 site which contained four replicates for all genotypes collected in the 2019 dataset. Separate analyses were conducted for each of the rating methods (5 raters, Ground Truth (GT), and the Machine Learning (ML) scores).

The data was analyzed with a mixed linear model to estimate variance components to calculate heritability and BLUPs. This model is based on the work of Gilmour, 1997. The response variable (Y) is obtained by taking the average of 3-46 (median of 27) plant observations (images) to generate a single value per EU as per protocol for measuring this trait.

Y = μ + G + Row + Col + Residual, where

Y = is the plant level scores averaged for each EU
μ = overall mean of dataset (fixed effect)
G = genotype (random effect)
Row, Col = row & column effects (random effect)
Residual = random error with an AR1 x AR1 structure (auto-regressive 1)

Broad sense heritability was calculated as H2 = Var(G) / (Var(G) + Var(residual)) and BLUPs for each genotype (G) were predicted. Median BLUP correlations were calculated as the Pearson correlation of the different BLUP rating methods (5 raters and ML) to the GT BLUPs and the mean BLUP score was calculated for each rating method. Scatterplots of all the rating methods were also generated.

## Results

### Phenotyping and human scoring

Phenotyping canola for blackleg severity requires screening for symptoms present in the crown-root interface during the period between seed-set and maturity. In this study, single-row plots of canola were grown across Canada in two growing seasons (2018 and 2019), and these plots were inoculated with blackleg inoculum when the majority of plants were at the 2 to 4-leaf stage. Plants were processed by cutting and imaging during the optimal window between seed-set and maturity. For each sample, two images were taken, one showing a cross-section view of the plant stem and another showing the side view of the stem. Approximately 25 plants were processed per plot. This process was repeated across the two growing seasons.

Blackleg severity was quantified using the ‘*Phoma Lingam* Crown Disease Severity/Score’ (PLCRDS), which is a 1 –9 scale with 1 denoting high disease severity and 9 denoting absence of disease, and is traditionally measured by experts trained to score samples based on severity of disease by looking at cross-sections of infected plants. Additionally, a rating of “Unsure” was created in cases where images could not be evaluated. For this study, five human experts scored images for all samples. Ground Truth (GT) estimates are represented by the median rating from all raters (see methods for details). GT data collected in 2018 and 2019 showed considerably different distributions, with 2018 data having the highest number of samples rated 9 (no disease) and 2019 data showing a depletion of scores of 3, 4, 5, and 9 (Figure 5).

**Figure 5:**
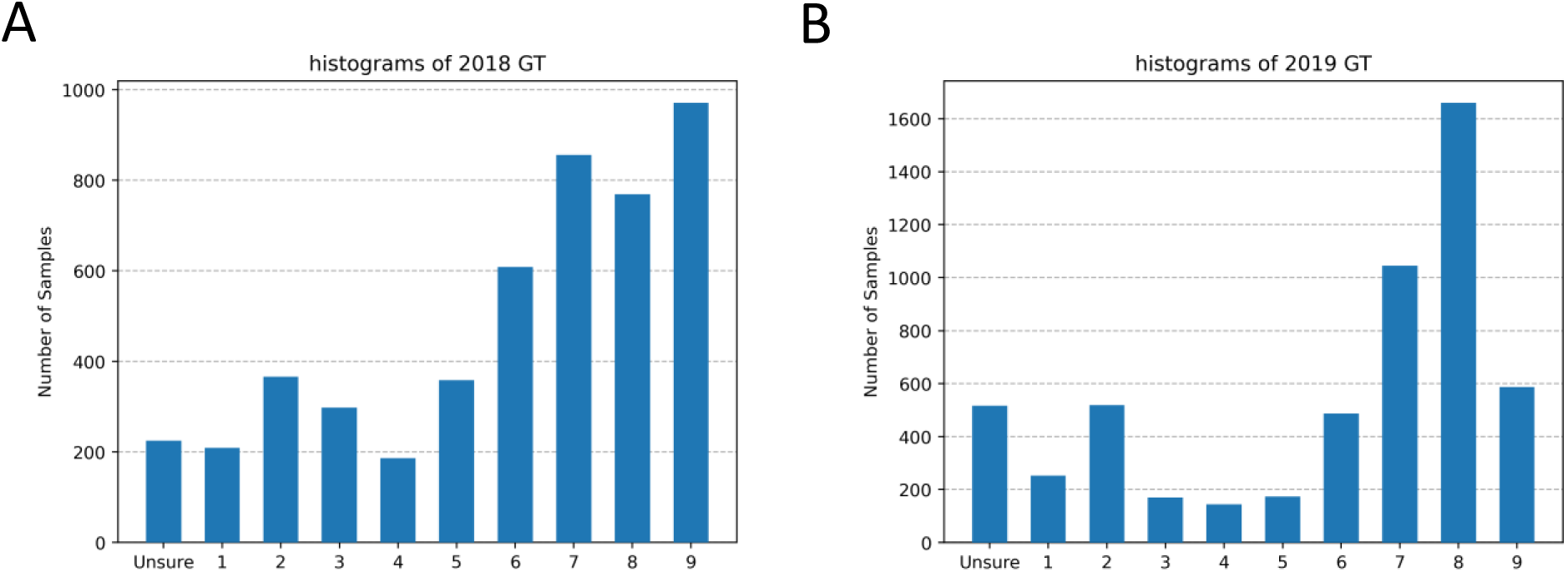
Histograms of Blackleg GT scores derived from 5 expert raters in 2018 (A) and 2019 (B) data. Low scores denote high susceptibility to Blackleg and high scores denote low susceptibility.

While expert raters are the gold standard for scoring, there is substantial variation between scores rated by different scorers (Table 1). Two metrics are used to indicate the performance and consistency of raters within years, across years, and between raters, and also to measure our method’s performance mentioned in later sections. One is accuracy, which is a standard metric that represents the percentage of the instances where one rater’s scores exactly matched GT ratings. The second is a customized accuracy metric, referred to as one-off accuracy (acc_oneoff), which represents the percentage of instances where one rater’s scores are within 1 scale of GT ratings. For one-off accuracy, since Unsure is a nominal category, all Unsure GT samples are excluded, and all Unsure observations are considered errors. Using these metrics, each rater’s performance against the GT within each year is calculated and they are ranked from best (rater 1) to worst (rater 5) based on their accuracy. Human raters varied with respect to the GT (Table 1 A-B). Also, while the highest ranked had similar metrics across years, the 3^rd^ through 5^th^ best raters all performed less accurately by both metrics in 2019 compared to 2018 (Table 1).

**Table 1:** Expert rater analysis. Variation in expert rater accuracy and accuracy within one label off when measured against the ground truth rating or other raters. (A-B) Accuracy and one-off accuracy against ground truth for 2018 (A) and 2019 (B) data. (C-F) Accuracy measured as consistency across raters or against the ground truth (GT). Accuracy (C, E) and one-off accuracy (D, F) is shown, and 2018 data is shown in C-D while 2019 data is shown in E-F. Note that in tables D and F, one-off accuracy scores for reciprocal comparisons may not be identical due to the treatment of Unsure ratings.

| 2018 Data | rater1 | rater2 | rater3 | rater4 | rater5 |
| --- | --- | --- | --- | --- | --- |
| accuracy | 0.719 | 0.680 | 0.650 | 0.624 | 0.557 |
| One-off_accuracy | 0.935 | 0.926 | 0.937 | 0.898 | 0.863 |

**B**
| 2019 Data | rater1 | rater2 | rater3 | rater4 | rater5 |
| --- | --- | --- | --- | --- | --- |
| accuracy | 0.741 | 0.689 | 0.536 | 0.531 | 0.291 |
| One-off_accuracy | 0.929 | 0.931 | 0.885 | 0.795 | 0.547 |

**C** 2018 raters Accuracy
|  | rater1 | rater2 | rater3 | rater4 | rater5 | GT |
| --- | --- | --- | --- | --- | --- | --- |
| rater1 | 1 | 0.517 | 0.474 | 0.486 | 0.458 | 0.719 |
| rater2 | 0.517 | 1 | 0.516 | 0.463 | 0.506 | 0.680 |
| rater3 | 0.474 | 0.516 | 1 | 0.551 | 0.388 | 0.650 |
| rater4 | 0.486 | 0.463 | 0.551 | 1 | 0.386 | 0.624 |
| rater5 | 0.458 | 0.506 | 0.388 | 0.386 | 1 | 0.557 |
| GT | 0.719 | 0.68 | 0.65 | 0.624 | 0.557 | 1 |

**D** 2018 raters one-off Accuracy
|  | rater1 | rater2 | rater3 | rater4 | rater5 | GT |
| --- | --- | --- | --- | --- | --- | --- |
| rater1 | 1 | 0.879 | 0.880 | 0.854 | 0.872 | 0.964 |
| rater2 | 0.879 | 1 | 0.871 | 0.838 | 0.832 | 0.954 |
| rater3 | 0.858 | 0.850 | 1 | 0.895 | 0.723 | 0.942 |
| rater4 | 0.846 | 0.831 | 0.91 | 1 | 0.744 | 0.918 |
| rater5 | 0.857 | 0.818 | 0.729 | 0.738 | 1 | 0.875 |
| GT | 0.935 | 0.926 | 0.937 | 0.898 | 0.863 | 1 |

**E** 2019 raters Accuracy
|  | rater1 | rater2 | rater3 | rater4 | rater5 | GT |
| --- | --- | --- | --- | --- | --- | --- |
| rater1 | 1 | 0.535 | 0.479 | 0.395 | 0.235 | 0.741 |
| rater2 | 0.535 | 1 | 0.521 | 0.384 | 0.232 | 0.689 |
| rater3 | 0.479 | 0.521 | 1 | 0.371 | 0.189 | 0.536 |
| rater4 | 0.395 | 0.384 | 0.371 | 1 | 0.368 | 0.531 |
| rater5 | 0.235 | 0.232 | 0.189 | 0.368 | 1 | 0.291 |
| GT | 0.741 | 0.689 | 0.536 | 0.531 | 0.291 | 1 |

Table 1: Variation in expert rater accuracy and accuracy within one label off when measured against the ground truth rating or other raters.
|  | rater1 | rater2 | rater3 | rater4 | rater5 | GT |
| --- | --- | --- | --- | --- | --- | --- |
| rater1 | 1 | 0.897 | 0.856 | 0.718 | 0.470 | 0.962 |
| rater2 | 0.876 | 1 | 0.903 | 0.678 | 0.440 | 0.942 |
| rater3 | 0.838 | 0.904 | 1 | 0.621 | 0.358 | 0.898 |
| rater4 | 0.723 | 0.699 | 0.639 | 1 | 0.701 | 0.83 |
| rater5 | 0.454 | 0.436 | 0.354 | 0.673 | 1 | 0.548 |
| GT | 0.929 | 0.931 | 0.885 | 0.795 | 0.547 | 1 |

Rater-to-rater comparisons were also performed in 2018 and 2019 by looking at accuracy and one-off accuracy (Table 1 C-F). These tables show that individually raters often do not agree with each other. The highest accuracy rating when comparing two reviewers was 0.551 between rater 3 and rater 4 in 2018. The majority of accuracy ratings between expert raters are below 0.5.

Finally, to investigate the consistency of human raters across time, a subset of the 2018 data (6%) was rated again in 2019 by 4 raters with two returning raters from 2018. The 4 raters’ ratings are used to create a GT referred to here as GT from 2019, in the same way that the GT generated based on ratings of 5 raters from 2018, which is referred to here as GT from 2018. We compare the difference between these two GTs by calculating the metrics of GT from 2019 against GT from 2018. For the two returning raters, year-to-year comparison is also calculated in the same way, such that raters’ ratings from 2019 are scored against their ratings from 2018. For the year-to-year comparison an individual rater’s accuracy ranges from 54-60% and one-off accuracy ranges from 86-87% (Table 2). Comparatively, GT has higher year-to-year consistency as the accuracy is 0.710 and one-off accuracy is 0.960. In conclusion, these analyses demonstrate that individual human experts exhibit a certain level of inconsistency and variance (hence errors), and the aggregated ratings of a group of human experts provide more consistent and accurate ratings. This also implies the need for an independent objective AI-based solutions that could help to provide a third-opinion or reference for the rating.

**Table 2:** Expert raters’ year-to-year comparison. Expert raters’ year-to-year comparison. Two returning raters evaluated the same images in 2018 and 2019, and their consistency across years was evaluated. The GT created by aggregating multiple raters rating from different years shows high consistency, while individual rater shows lower consistency of the ratings made from different years.

| **year-to-year**<br>**(2019 against**<br>**2018)** | **GT** | **Returning**<br>**rater 1** | **Returning**<br>**rater 2** |
| --- | --- | --- | --- |
| **Accuracy** | **0.710** | **0.540** | **0.601** |
| **One-Off Accuracy** | **0.960** | **0.871** | **0.863** |

### Image-level evaluation and model performance

The two-stage models’ performance was evaluated by treating each pair of images as an individual sample. The first-stage model is evaluated standalone using the 2018 testing set. Due to the relative simplicity of the task for this model in detecting if there is a valid cross-section or not, the metrics for the model’s performance is the accuracy of finding a cross-section or not versus the ground-truth of whether there is a cross-section or not. The accuracy on the 2018 testing set reaches 99.613% (12 errors out of 3097 samples). These errors are manifested in 3 types: multiple bounding box (BB) detection with high confidence scores above the threshold, false BB detection on Unsure samples, and miss a possible cross-section. Examples of the correct detections are in Figure 3, and examples of errors are in Figure 4.

For the second-stage model, the output of the first-stage model is used as input for scoring, and the output scores are compared to the GT based on the human raters’ ratings. The same metrics of accuracy and one-off accuracy mentioned above are used here to measure the model’s performance. For the model’s results, different versions are generated by using several different second-stage confidence thresholds based on our empirical judgement. The evaluation is broken down on testing set 2018 and 2019 and compared to raters sorted by their performance from rater1 to rater5 (See Table 3). The evaluation shows that the model’s performance is slightly above the median level of human raters (rater3), but below the first two best human raters, based on both metrics on both years’ testing sets. It also indicates that the threshold value has minimum impact on the model’s performance as long as it is within the proper range. For simplicity, for any model related analysis below, one of the best versions by the 0.25 threshold is used.

**Table 3:** Model performance. Performance of the proposed model with different second-stage thresholds compared with the 5 expert raters’ ratings against the GT on 2018 (A) and 2019 (B) testing sets.

| 2018 Test set | rater1 | rater2 | rater3 | rater4 | rater5 | Model Result |  |  |
| --- | --- | --- | --- | --- | --- | --- | --- | --- |
|  |  |  |  |  |  | Stage-2<br>th=0.30 | Stage-2<br>th=0.25* | Stage-2<br>th=0.20 |
| Accuracy | 0.732 | 0.717 | 0.679 | 0.640 | 0.578 | 0.701 | 0.698 | 0.695 |
| Acc_oneoff | 0.940 | 0.938 | 0.942 | 0.907 | 0.875 | 0.926 | 0.923 | 0.921 |

Table 3: Performance of the proposed model with different second-stage thresholds compared with the 5 expert raters' ratings against the GT on 2018 (A) and 2019 (B) testing sets.
| 2019 Test set | rater1 | rater2 | rater3 | rater4 | rater5 | Model Result |  |  |
| --- | --- | --- | --- | --- | --- | --- | --- | --- |
|  |  |  |  |  |  | Stage-2<br>th=0.30 | Stage-2<br>th=0.25* | Stage-2<br>th=0.20 |
| Accuracy | 0.741 | 0.689 | 0.536 | 0.531 | 0.291 | 0.604 | 0.607 | 0.607 |
| Acc_oneoff | 0.931 | 0.933 | 0.894 | 0.810 | 0.579 | 0.907 | 0.919 | 0.921 |

To further investigate the model’s performance, it is important to note that this model is heavily trained on 2018 data and only on 1% of 2019 data, and the model is tested on the remaining 99% of 2019 collection. For an ablation study about the data’s impact on the model’s performance, a model of the same structure is trained on only the same 2018 training set and tested on the same 2018 and 2019 testing sets as the proposed model. The performance of the 2018-only trained model and the proposed model, and the median and 4^th^ best rater are in Table 4. The table shows that the 2018-only trained model performs similarly to the proposed model, and to the median rater on 2018 data. However, it performs worse than the proposed model and the median rater on 2019 data. This might be because 2019 data could be more difficult to rate as revealed by the lower accuracy of human raters on 2019 data (or larger variance). From the opposite perspective, this means that adding only 1% of 2019 data in the model training helps the model’s generalization and stability. This implies that for future data, with the same imaging system, the proposed model would likely be above the level of 2018- only-trained model on 2019 testing set, and potentially equivalent to its performance on 2019 testing set, which is slightly above the median level of median raters. Therefore, the proposed model achieves performance within the accuracy range of median human raters with the image-wise evaluation.

**Table 4:** 2018-only trained model. Performance of 2018-only trained model and the proposed model compared with rater3 and rater4 on 2018 and 2019 testing sets, and their year-to-year differences. Note that rater3 and rater4 for the two years might not be the same rater since these are alias sorted by rater accuracy of each year.

| Accuracy/One-off Accuracy | On 2018 testing set | On 2019 testing set | Change in absolute value |
| --- | --- | --- | --- |
| 2018-only trained model | 0.687/0.933 | 0.439/0.845 | -0.248/-0.088 |
| The proposed model | 0.698/0.923 | 0.607/0.919 | -0.091/-0.004 |
| rater3 | 0.679/0.942 | 0.536/0.894 | -0.143/-0.048 |
| rater4 | 0.640/0.907 | 0.531/0.810 | -0.109/-0.097 |

### Heritability of blackleg severity determined by machine learning

To further validate the model, broad sense heritability was calculated for all rating methods (5 raters, Ground Truth (GT), and the Machine Learning (ML) scores) using data generated in 2019 (Table 5). One location (Ardrossan, AB) was chosen for analysis as it contains 4 replicates of all genotypes used in the study. The heritability estimates were calculated from a linear mixed model with spatial adjustments. These results show that ML auto-scoring rating achieves a similar heritability estimate as to the expert raters and the ground truth estimate.

**Table 5:** Heritability across rating methods. Heritability estimates are similar for Canola Blackleg scores for human raters, ground truth (GT) and machine learning (ML) ratings. Columns show the genotype variance (GE_var), residual (resid), and heritability (H2) estimates.

| trait | GE_Var | resid | H2 |
| --- | --- | --- | --- |
| ML | 0.19 | 0.30 | 0.38 |
| Rater_1 | 0.26 | 0.38 | 0.41 |
| Rater_2 | 0.35 | 0.35 | 0.51 |
| Rater_3 | 0.36 | 0.40 | 0.48 |
| Rater_4 | 0.26 | 0.56 | 0.31 |
| Rater_5 | 0.18 | 0.24 | 0.44 |
| GT | 0.31 | 0.37 | 0.46 |

The linear mixed model was then used to calculate BLUPs for each genotype for each rating method (Table 6). The overall mean score of the ML BLUPs (7.49) is close to the overall mean score across all 5 expert raters (7.05). The expected BLUP variation for all rating methods is seen among the different genotypes with the lowest score seen for the genotype WESTAR, which is the highly blackleg susceptible control. Even though there is a mean shift in ratings based on the rating method, the rankings between all methods are very similar. The ML method shows a similar re-ranking as is seen between the expert raters. The variation among the expert raters BLUPs is similar to the variation between the auto-scoring and expert raters (Figure 6). Overall, the BLUPs are well correlated between all rating methods.

**Figure 6:**
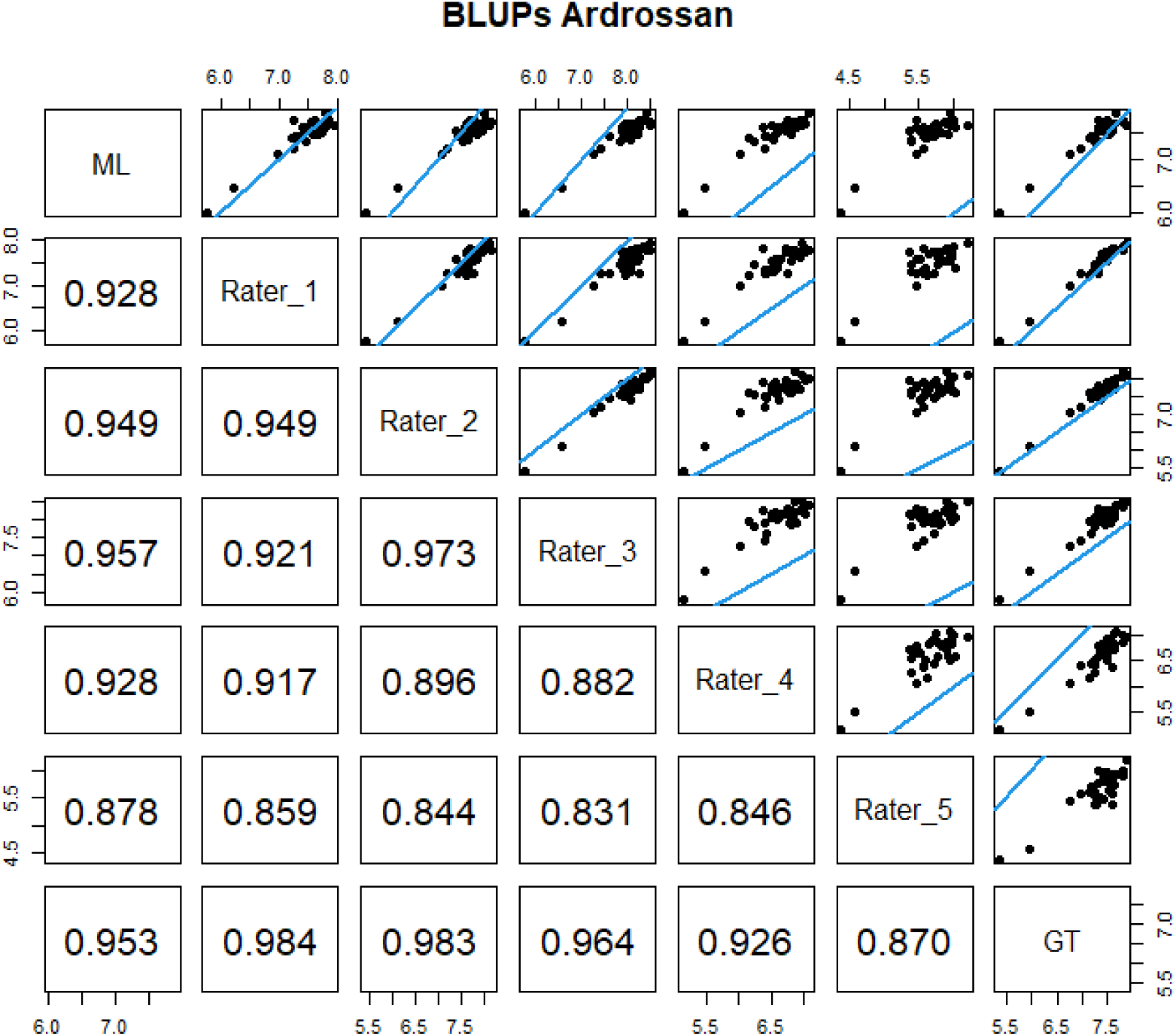
Numerical correlation values and dot plots of Best Linear Unbiased Predictions (BLUPs) for machine learning (ML), expert raters 1-5, and ground truth (GT) values. The blue line shows the equality line (x=y).

**Table 6:** BLUPs across rating methods and genotypes. The Best Linear Unbiased Predictions (BLUPs) for each genotype are shown for expert raters and machine learning ratings. Each row shows an individual genotype BLUP, the median correlation, or the mean score.

| Genotype | ML | Rater_1 | Rater_2 | Rater_3 | Rater_4 | Rater_5 | GT |
| --- | --- | --- | --- | --- | --- | --- | --- |
| PHBL001 | 7.64 | 7.94 | 8.13 | 8.47 | 6.97 | 6.19 | 7.85 |
| PHBL002 | 7.74 | 7.87 | 8.06 | 8.34 | 7.00 | 5.99 | 7.80 |
| PHBL003 | 7.70 | 7.77 | 8.19 | 8.49 | 6.85 | 5.89 | 7.78 |
| PHBL004 | 7.88 | 7.79 | 8.01 | 8.39 | 7.08 | 5.94 | 7.64 |
| PHBL005 | 7.69 | 7.74 | 7.94 | 8.14 | 7.03 | 5.74 | 7.62 |
| PHBL006 | 7.67 | 7.82 | 7.71 | 8.17 | 6.73 | 5.90 | 7.62 |
| PHBL007 | 7.70 | 7.79 | 7.87 | 8.05 | 6.82 | 5.94 | 7.61 |
| PHBL008 | 7.61 | 7.80 | 7.88 | 8.25 | 6.37 | 5.58 | 7.60 |
| PHBL009 | 7.55 | 7.74 | 7.97 | 8.16 | 6.72 | 5.37 | 7.58 |
| PHBL010 | 7.61 | 7.75 | 7.84 | 7.89 | 6.90 | 5.69 | 7.54 |
| PHBL011 | 7.64 | 7.62 | 7.80 | 8.10 | 6.83 | 5.78 | 7.53 |
| PHBL012 | 7.47 | 7.68 | 7.71 | 8.04 | 6.80 | 5.65 | 7.51 |
| PHBL013 | 7.64 | 7.53 | 7.80 | 8.13 | 6.50 | 5.95 | 7.49 |
| PHBL014 | 7.77 | 7.63 | 7.71 | 8.24 | 6.93 | 5.98 | 7.49 |
| PHBL015 | 7.50 | 7.58 | 7.88 | 8.20 | 6.57 | 5.92 | 7.49 |
| PHBL016 | 7.55 | 7.55 | 7.63 | 7.87 | 6.82 | 5.80 | 7.44 |
| PHBL017 | 7.51 | 7.74 | 7.62 | 7.91 | 6.78 | 5.81 | 7.43 |
| PHBL018 | 7.62 | 7.58 | 7.77 | 8.12 | 6.66 | 5.47 | 7.41 |
| PHBL019 | 7.71 | 7.55 | 7.72 | 8.23 | 6.82 | 5.88 | 7.41 |
| PHBL020 | 7.69 | 7.68 | 7.64 | 7.96 | 6.82 | 5.97 | 7.39 |
| PHBL021 | 7.56 | 7.58 | 7.41 | 8.05 | 6.63 | 5.50 | 7.36 |
| PHBL022 | 7.55 | 7.47 | 7.58 | 8.02 | 6.59 | 5.76 | 7.31 |
| PHBL023 | 7.77 | 7.27 | 7.78 | 8.27 | 6.76 | 5.47 | 7.31 |
| PHBL024 | 7.63 | 7.39 | 7.59 | 8.06 | 6.58 | 6.01 | 7.30 |
| PHBL025 | 7.39 | 7.31 | 7.66 | 8.08 | 6.54 | 5.40 | 7.27 |
| PHBL026 | 7.40 | 7.36 | 7.66 | 8.06 | 6.51 | 5.60 | 7.25 |
| PHBL027 | 7.35 | 7.48 | 7.54 | 7.82 | 6.25 | 5.39 | 7.24 |
| PHBL028 | 7.44 | 7.35 | 7.52 | 7.92 | 6.40 | 5.56 | 7.22 |
| PHBL029 | 7.43 | 7.28 | 7.45 | 7.62 | 6.42 | 5.72 | 7.16 |
| PHBL030 | 7.41 | 7.23 | 7.63 | 7.94 | 6.16 | 5.62 | 7.13 |
| PHBL031 | 7.20 | 7.26 | 7.21 | 7.42 | 6.40 | 5.57 | 6.97 |
| PHBL032 | 7.11 | 6.99 | 7.07 | 7.26 | 6.04 | 5.45 | 6.74 |
| PHBL033 | 6.46 | 6.22 | 6.12 | 6.58 | 5.48 | 4.58 | 5.97 |
| WESTAR | 6.01 | 5.77 | 5.43 | 5.77 | 5.15 | 4.39 | 5.37 |
| Median correlation | 0.95 | 0.98 | 0.98 | 0.96 | 0.93 | 0.87 | 1.00 |
| Mean Score | 7.49 | 7.47 | 7.60 | 7.94 | 6.59 | 5.66 | 7.32 |

## Discussion

Canola blackleg is a leading cause of Canola yield loss worldwide (Zheng et al., 2020). In addition to fungicides and management practices, planting resistant varieties has been highly effective in reducing yield loss. Blackleg susceptibility is highly variable across genotypes yet can be difficult to score accurately and consistently. Accurate scoring for blackleg is important for several key reasons. From a regulatory standpoint, accurate scoring for all registered varieties grown in Canada helps to mitigate the economic impacts of blackleg infection. Additionally, plant breeders make breeding and advancement decisions about resistant varieties based on severity scoring in test plots. Severity scoring in test plots has the additional complication of field and environmental variability, so it is beneficial to reduce variability in scoring as much as possible for robust predictions. Accurate scoring can also aid in identification and cloning of resistance genes, opening the door to genome edited blackleg disease resistance. Taken together, accurate scoring of blackleg infection is key to improving desired outcomes.

While a team of expert raters is the gold standard for scoring blackleg disease, human raters can be inconsistent, expensive, and logistically complicated to implement during the short time window when blackleg severity can be scored. Furthermore, the human raters need substantial training to understand and implement a rating scale. While there has been progress in creating a single scoring rubric and set of training documents, scores are still assigned subjectively based on human interpretations of the reference images. In contrast, our machine learning model is as accurate on average as the median expert rater, and since it is pre-trained, it can be consistently implemented by any field team regardless of experience. Utilizing a machine learning algorithm can circumvent the need for highly trained field staff, saving time and money while maintaining the high standards required for accurate blackleg severity scoring.

The results shown above do exhibit year-to-year differences, both for the model and expert raters. The model performs better on 2018 data, but worse on 2019 data. The year-to-year difference of the model might be due to several factors. First, the difference of the data in 2018 and 2019 would affect the model’s performance. To illustrate that, the histograms of 2018 and 2019 data GT are in Figure 5, which shows that there is a difference in the score distribution between the two years’ data. While neither year has perfectly balanced data, 2018 data is more evenly distributed, and 2019 data is more unimodal with the highest number of samples scoring an 8. In addition, human raters in general perform worse on 2019 set compared to 2018 (Table 1), which implies that 2019 data might have more ambiguous samples due to lower overall disease pressure, or the raters rating on 2019 data might be less experienced in general, hence might yield less reliable or consistent ratings.

In addition to the current application of machine learning for blackleg severity scoring, there are several future opportunities that will likely improve the accuracy and efficiency of auto-scoring. For example, mobile phone cameras may be used in place of the specific cameras in this study, improving access and decreasing cost of auto-scoring. This is particularly important in large canola-growing regions with limited access to trained raters since an individual team of experts has limits on how many fields they can score within the narrow window needed for accurate scoring. Furthermore, multi-spectral or hyperspectral sensors may be used in place of RGB cameras, increasing the ability to detect subtle changes in disease progression.

In conclusion, here we demonstrate an accurate and reliable method for canola blackleg severity scoring using machine-learning enabled image analysis. This method is as accurate as the median rater at scoring disease severity and produces similar heritability and BLUP estimates as expert raters. Future work to improve hardware and software may result in cheaper, more accurate blackleg scoring that can be implemented at a larger scale.

## Data Availability Statement

The datasets presented in this article are proprietary to Corteva Agrisciences. Requests to access the datasets should be directed to Daniel Stanton.

---

**Supplemental Figure 1.**
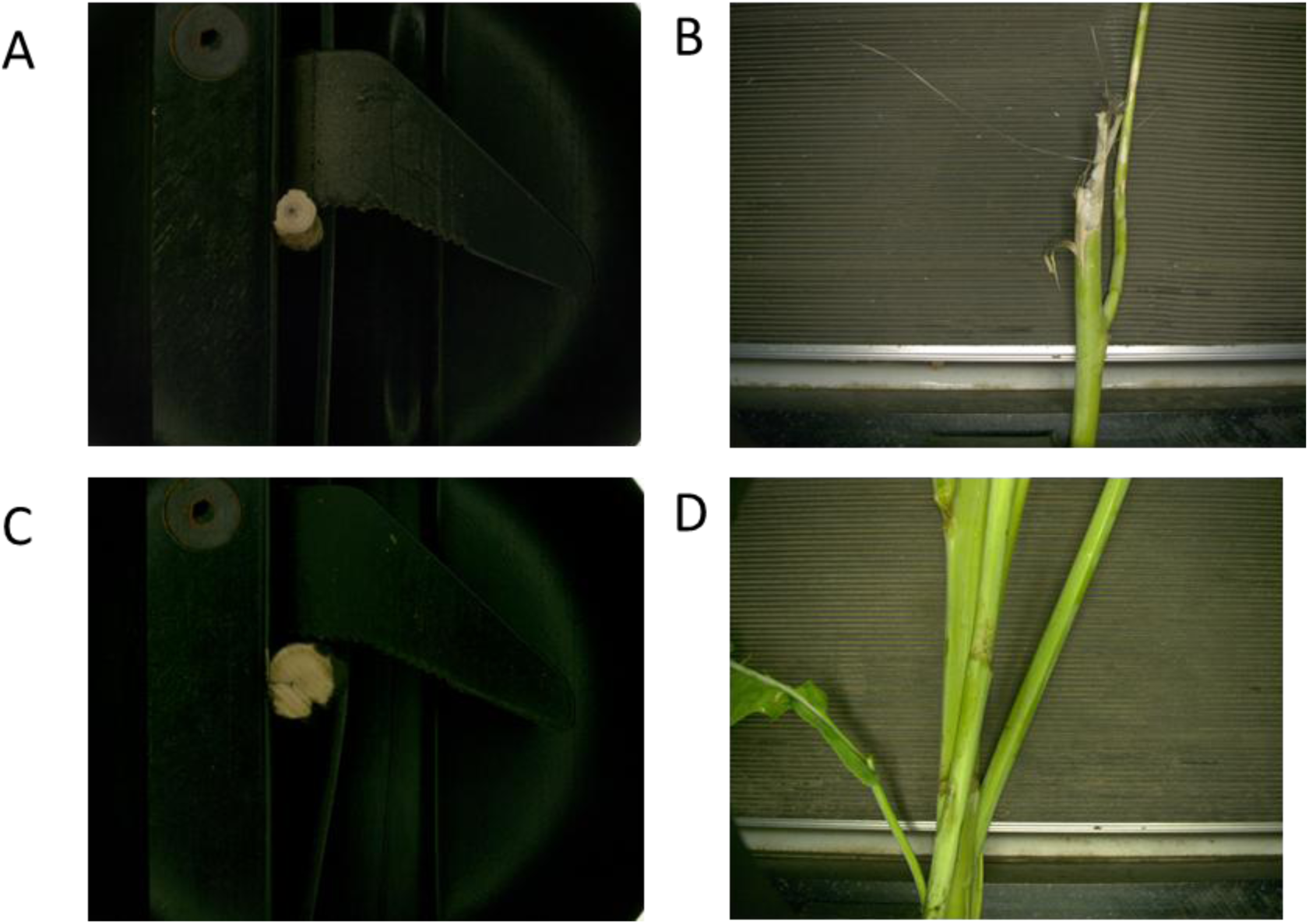
Examples of paired images collected in 2018 (A-B) and 2019 (C-D).

**Supplemental Table 1.**
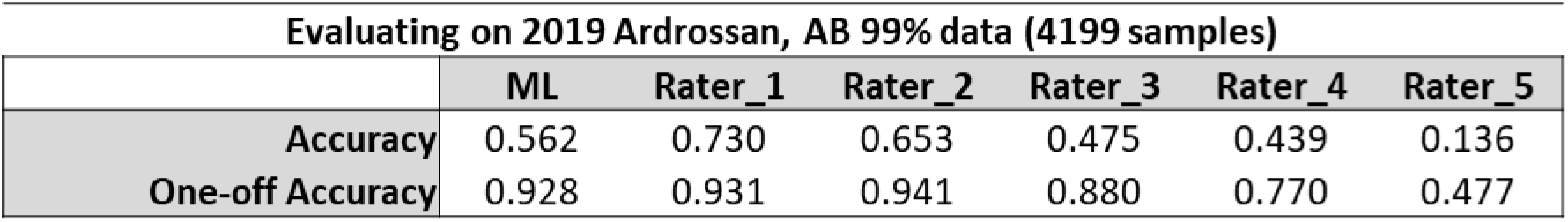
Comparison of expert rater and machine learning model accuracy on the subset of data from Ardrossan, AB used for the heritability analysis. Accuracy and accuracy within one label off are calculated vs the GT.

